# Validation and standardization of a new radial-arm water maze protocol using a murine model of mild closed head traumatic brain injury

**DOI:** 10.1101/2020.04.24.059329

**Authors:** Teresa Macheda, Kelly N. Roberts, Adam D. Bachstetter

## Abstract

Cognitive impairments can be a significant problem after a traumatic brain injury (TBI), which affects millions worldwide each year. There is a need for establish reproducible cognitive assays in rodents to better understand disease mechanisms and to develop therapeutic interventions towards treating TBI-induced impairments. Our goal was to validate and standardize the radial arm water maze (RAWM) test as an assay to screen for cognitive impairments caused by TBI. RAWM is a visuo-spatial learning test, originally designed for use with rats, and later adapted for mice. The present study investigates whether test procedures, such us the presence of extra-maze cues influences learning and memory performance. C57BL/6 mice were tested in an 8-arm RAWM using a four-day protocol. We demonstrated that two days of training, exposing the mice to extra-maze cues and a visible platform, influenced learning and memory performance. Mice that did not receive training performed poorer compared to mice trained. To further validate our RAWM protocol, we used scopolamine. We, also, demonstrated that a single mild closed head injury (CHI) caused deficits in this task at two weeks post-CHI. Our data supported the use of 7 trials per day and a spaced training protocol as key factor to unmask memory impairment following CHI. Here, we provide a detailed standard operating procedure for RAWM test, which can be applied to a variety of mouse models including neurodegenerative diseases and pathology, as well as when pharmacological approaches are used.

## Introduction

Traumatic brain injury (TBI) is a major public health problem worldwide and leads to temporary or permanent physical and cognitive impairments. In particular, people with a history of TBI have an increase risk to develop dementia and neurodegenerative disease [1,2].

Mechanisms of selective vulnerability to cognitive deficits following a mild TBI are still not well understood, and the use of animal models accelerate a better understanding of the pathological and behavioral outcome associated with a mild TBI. Novel object recognition (NOR) and Morris water maze (MWM) test are the two most popular assays used to evaluate cognitive function after mild TBI [3]. While the radial arm water maze (RAWM) task is becoming a standard tool to assess memory in rodents, only a few studies have used it as tool to evaluate memory in mice after mild TBI [3], and no study has validated optimal methods for the behavior following a mild TBI.

In 1984, Buresova et al. described a “radial maze in the water tank”, making this the first time that water was used as an aversive stimulus in a radial maze [4]. A significant advantage of RAWM is that food deprivation is not required, and odors that could be used by the animal as cues are eliminated. A few years later, Hyde et al. [5] tested three inbred strains of mice in RAWM. Since then, several protocols and multiple variations (4-12 arms) of the maze have been used, with RAWM protocols as short as 2 days, where mice undergo up to a total of 30 trials [6] or 20 trials [7]. Other protocols, instead, are 7 days long and mice are trained in four trials per day [8]. These are just a few examples of a large variety of RAWM protocols used in many different laboratories.

Training schedule affects level of acquisition in humans [9], rodents [10–16], invertebrates such as the mollusk *Aplysia California* [17–20], *Drosophila Melanogaster* [21], bees [22], and in non-human primates [23]. In particular, it has been shown that a more robust memory formation is linked to spaced training rather than a massed training [10–13, 24, 25] where short or no intervals are employed. Moreover, irregular spaced training may enhance learning [26].

The goal of this paper is to test if mice trained in a spaced RAWM training protocol have the ability to learn the task with a total of 28 trials spaced over 4 days. We hypothesized that reducing the number of trials per day would improve their ability to learn and reduce fatigue. An added benefit is the ability to increase the capability to test more mice in one session. The result of our study, support our hypothesis that few trials per day spaced over 4 days, is sensitive at detecting cognitive deficits following a mild closed head injury (CHI). Our RAWM protocol will likely be widely applicable to detect cognitive deficits in other mouse models of injury or disease.

## Materials and Methods

### Chemical

Scopolamine hydrobromide was purchased from Sigma (cat. no.6533-68-2). It was dissolved in double-distilled water, sterile filtered (0.2 um sterile filter; VWR North America, cat. # 76012-774) and administered at a dose of 1 mg/Kg/10 ml. Intraperitoneal (ip) injections were performed 30 minutes before trial 1 every day during the 4-day test. Scopolamine was freshly prepared each day animals were dosed.

### Animals

All animal procedures were approved by the Institutional Animal Care and Use Committee (IACUC) of the University of Kentucky and experiments were conducted in accordance with the standards of proper experimentation in the Guide for the Care and Use of Laboratory Animals and ARRIVE guidelines.

The study used 141 adult mice (72/69 ♀/♂) 3-4 months old, C57BL/6J mice (Jackson laboratory, Bar Harbor, ME; stock number: 000664). The number of mice used for each experiment is reported in the figure legend. Animals were group-housed (4-5 per cage) in a controlled humidity (43-47%) and temperature (22-23° C) environment and 12/12-h (7 am-7 pm) light/dark cycle with free access to food and water. Behavioral experiments were conducted by the same operator (TM) and performed between 7.30 a.m. and 3.30 p.m.

Mice were assigned randomly to groups before the start of each experiment. Each cage had at least one animal from each group/ treatment in a random order and the person performing the experiments scopolamine injections (KNR) and RAWM test (TM) was blinded to treatment. Also, the person performing the RAWM test (TM), three days before the beginning of the experiment transferred the mice to a clean cage, marked the tail for easy identification, and handled them for the following three days. For the handling, a mouse was allowed to explore the experimenter’s arm and hand for 1-2 min, then returned to its home cage. Mice were never exposed to the maze before the start of the experiment. During the behavioral tests, mice were transferred to the experimental room and relocated in a clean recovery cage without bedding at least 30 minutes before the start of the test. The recovery cage had paper towels inside and was placed half on a heating pad to help the mice recovery from the swimming activity and half off the heat. Also, during the test, wet paper towels were promptly replaced, so mice could have a faster recovery in a dry and warm recovery cage. Male and female mice were tested in separate cohorts.

### Radial arm water maze apparatus

The RAWM test was performed in a circular pool (diameter=121 cm, depth 75 cm, Fig.1 A, B) (MazeEngineers, Boston, MA). A base (9 cm high), and an 8-shaped platform were added to the pool and eight identical metal inserts having a V shape (approximately 65 cm high by 42 cm long) were inserted in the pool to make an 8 arm RAWM (Fig.1 A, B). Arms were raised 9 cm above the level of water, to discourage mice from climbing over the inserts and jumping in the dead area of the pool or on the floor. The arms were made of stainless steel to avoid corrosion due to the continuous contact to the water. The water was made opaque by the addition of white liquid non-toxic paint (Sax 2684 Versatemp Non-Toxic Heavy Body Tempera Paint), and the water temperature was held constant at 21 ± 1° C. The escape platform was a circular (diameter 8 cm, 57 cm high, Fig.1 C) clear acrylic adjustable platform submerged 1 cm below the water level in one of the eight arms and was defined as the “goal arm”. The apparatus was isolated from the rest of the room by double black black-out curtains. Four extra-maze cues (triangle, square, checkerboard pattern and cross made of white corrugated plastic and black vinyl material; Fig.1 E) were hung on the inside of these curtains around the pool (Fig.1 A) at a height of 12 cm from the top of the pool to the cues. Four dimmable overhead lights were used and light intensity was kept between 4 and 6 lux in each arm and center of the maze. The platform was made visible by a flag (16 cm high, Fig.1 D) when needed.

**Figure 1:**
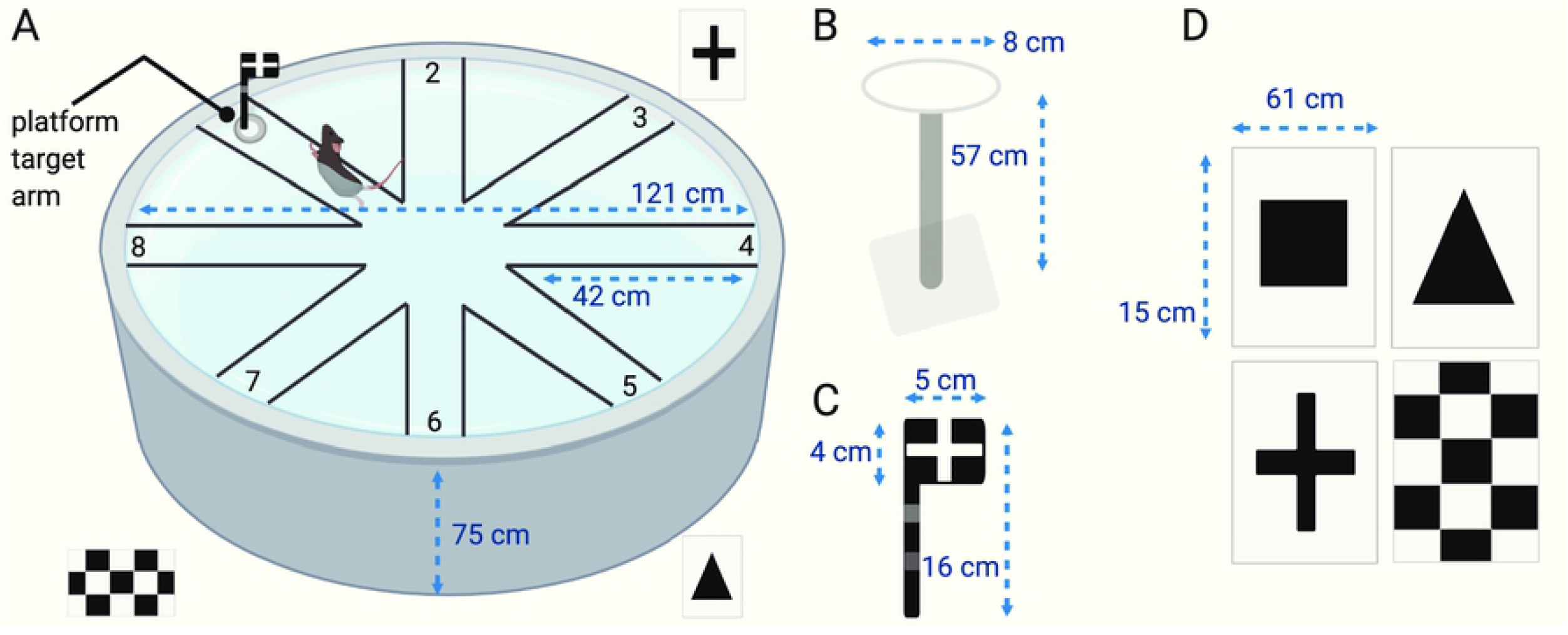
Pool and cues used for RAWM test. **(A)** RAWM pool is shown with a mouse swimming to reach the visible platform. Four cues (square-not shown in the picture-, triangle, cross and checker board) are equally spaced around pool. **(B)** Dimensions of platform, **(C)** flag and **(D)** cues are reported as reference.

A camera was positioned directly above the center of the pool and all experiments were recorded. EthoVision XT 11.0 (Noldus Information Technology) was used for video recording and scoring behavior.

### Radial Arm Water maze 7-trial protocol

Mice were tested for a total of 4 days and received 7-trial per day. To reduce fatigue, the 7 trials were divided in two blocks: *block 1*(*trials 1-3*) and *block 2* (*trials 4-7*). To encourage mice to learn the location of the platform, an alternation of visible and hidden platform was used during block 1 of training days (trial 1 and 3, day 1 and 2) and a hidden platform was used during block 2 (Fig. 2 A). During testing days, the platform was hidden in both block 1 and block 2 (Fig. 2 B).

**Figure 2:**
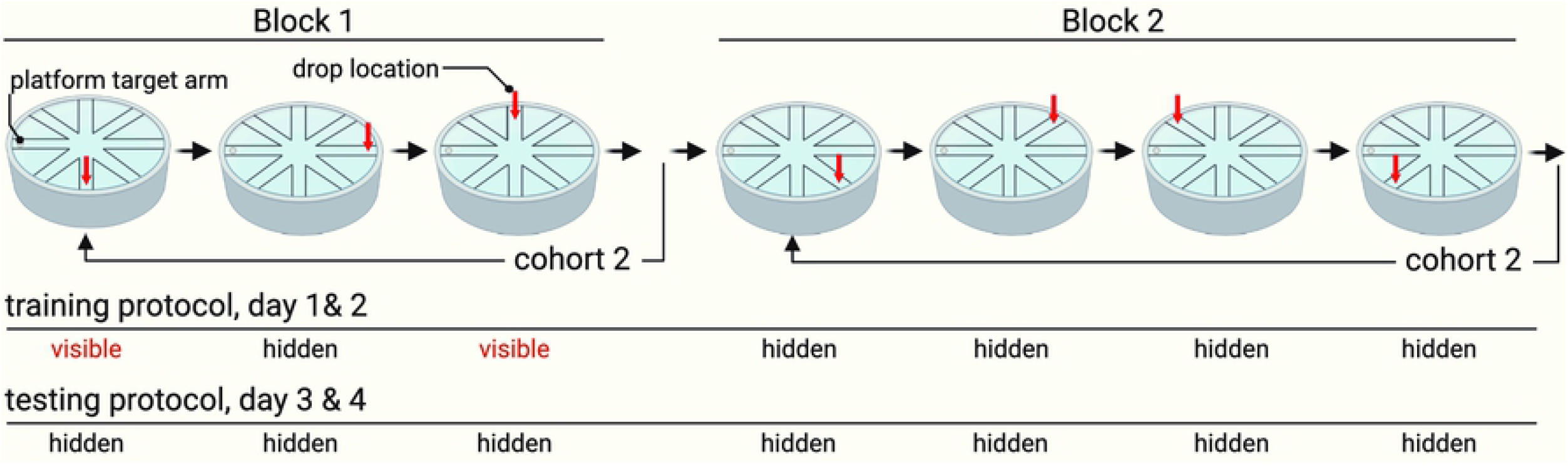
Step-by-Step procedure of RAWM test during training and testing days. During training days (day 1 and 2), the platform is made visible by a flag during block 1 (trial 1 and 3). During the testing days (day 3 and 4), the platform is hidden during all trials. The drop location of the mice varies in a semi-random fashion as shown by the red arrow. To reduce fatigue and learning limitation, a staggered design is used in RAWM test, with one cohort (10-15 mice) tested in a block before and then a second cohort is tested.

A staggered design was used with both cohort 1 and 2 completing block 1 before moving on to block 2 (Fig. 2 C). Mice that did not find the platform in 60 seconds were gently guided to it. Each mouse was given 15 seconds on the platform at the end of each trial to explore and memorize the spatial cues. When the trial was over, the mouse was gently removed from the pool, dried with a towel and returned to a heated drying recovery cage before the next trial. The goal arm was the same during all the trials and between mice, but the drop location varied between trials in a semi-random fashion (Fig. 2 A and B). To evaluate if the animals were using extra-maze cues to locate the platform, a group of mice was tested in the same maze but the extra-maze cues were removed and the platform was never made visible during the 4-day test. To reduce learning limitations due to fatigue and massive training [10–14, 24, 25] mice were tested in cohorts containing 10-15 mice each.

An error was scored every time the mouse entered an arm that did not contained the platform or when it entered the goal arm without escaping. If the mouse spent more than 15 seconds in the same zone, arm or center, it was also counted as an error. Total number of errors, latency to escape, distance and velocity were recorded.

A step-by-step standard operating procedure has been provided and it is available on www.protocols.io (https://www.protocols.io/edit/7-trail-rawm-bb84iryw/steps)

### Closed head injury surgery

The mild closed Head Injury (CHI) model was used in this study to demonstrate the applicability of this new behavioral protocol. CHI was performed as previously described using a digital stereotactically (Stoelting) guided electromagnetic impactor device to produce a highly reproducible TBI with minimal mortality [27, 28]. Mice were anesthetized with 5% isoflurane before the surgery and kept under anesthesia with continuous inhalation of isoflurane (2.5-3%, 1L/min) through a nose cone during surgery. Before the surgery body weight was recorded, the head was shaved, sterilized with 70% ethanol and 4% lidocaine cream was applied. Each mouse was secured in a digital stereotaxic frame (Stoelting; Wood Dale, IL, USA) using ear bars. The skull exposed after a midline sagittal incision was made. A 1 ml latex pipette bulb was placed under the head of the mouse and filled with water, this helped to diffuse the force of the impact away from the ear bars.

CHI mice received a single controlled midline impact (coordinates: mediolateral, 0.0 mm; anteroposterior, 1.5 mm) 1.0 mm deep with a controlled velocity of 5.0+0.2 m/s and a dwell time of 100 ms using a stereotaxic electromagnetic impactor (Impact one, Leica; Buffalo Grove, IL, USA) equipped with a 5.0 mm flat steel tip. Sham-injured mice received identical surgical procedures as the CHI group, but no impact was delivered. Following impact, the incision was sutured, body weight was monitored up to 5 days postsurgery. Sutures were removed 1-week post-surgery. Starting at 14 days post-surgery, mice were tested in RAWM test.

### Statistical Analysis

JMP Pro software version 14.0 (SAS institute, Cary, NC, USA) was used for statistical analysis. Graphs were generated using GraphPad Prism version 8.0. Summary values for the learning curves are expressed as mean ± SEM or median ± SEM, as noted in the figure legend. Boxplot were generated using the Tukey method. Scatter plots represent individual mice. Mouse numbers used are indicated in the figure. Differences were considered statistically significant when p < 0.05.

The median of errors made by the animal each day was considered for analysis. Only hidden trials were used in the analysis. The AUC was calculated using the trapezoid method, dividing the whole AUC into trapezoidal segments and counting the area of each segment separately, this was done for both errors and distance [29]. The sum of the area of all trapezoids is the total AUC, visible and hidden trials were used for the analyses. We used the following formula:

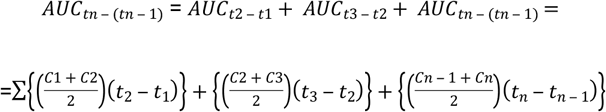

*Where:*

*AUC= area under the curve*
*c= errors or distance recorded at a specific time point*
*t_n_= time of observation of errors or distance at c_n_*
*t_n-1_= time of prior observation corresponding of errors or distance at c_n-1_*

A mixed model design using a standard least squares method was used to test for statistical differences between groups. Two models were compared. The first model included: day of testing, experimental group, sex of mice, and all the interactions. The second model did not include sex of mice (i.e., day of testing, experimental group and interaction). Overall, we did not find a statistically significant effect of sex on the outcome measures and did not find that the inclusion of sex improved our statistical model. Therefore, we report the results of the second statistical model without sex of the mice included in the model.

We have shown that the sex of the mice affects response to the CHI for some variables [30]; thus, the data for the CHI experiments were disaggregated by sex.

## Results

### Experiment 1: Use of cues in RAWM improves learning and memory

The goal of this experiment was to determine if cues were being used by the mice for the visuo-spatial learning during RAWM test. Mice were tested for a total of 4 days. We evaluated if the use of extra-maze cues and a visible escape platform is essential for the mice to learn the task and has a positive influence on RAWM output, or if the mice were using other extra maze cues or non-visuo-spatial search strategies. A 7-trial protocol was used to address this question, in particular two conditions were considered: 1) mice were exposed to the pool that was equipped with 4 extra-maze cues and a visible platform was used during training days (Fig. 2 A); 2) in this group of mice no extra-maze cues were used and mice were never allowed to use a visible platform, mice were never trained. As shown in Figure 3A, we found that there was a difference between mice trained (cues and visible platform) to locate the goal arm compared to the mice without cues or access to visible platform (p=0.0023). Also, the AUC (errors x trials) indicated that mice used cues to learn RAWM assay quicker compared to mice not exposed to cues (Fig. 3 B, p=0.0003). In fact, number of errors made by the animals using cues and a visible platform was significantly lower compared to mice not using any kind of cues. Finally, both the total distance travelled (Fig. 3 C, p=0.0031) and AUC (distance x trials) (Fig. 3 D, p=0.0007) were significantly higher in mice not exposed to the use of cues.

**Figure 3:**
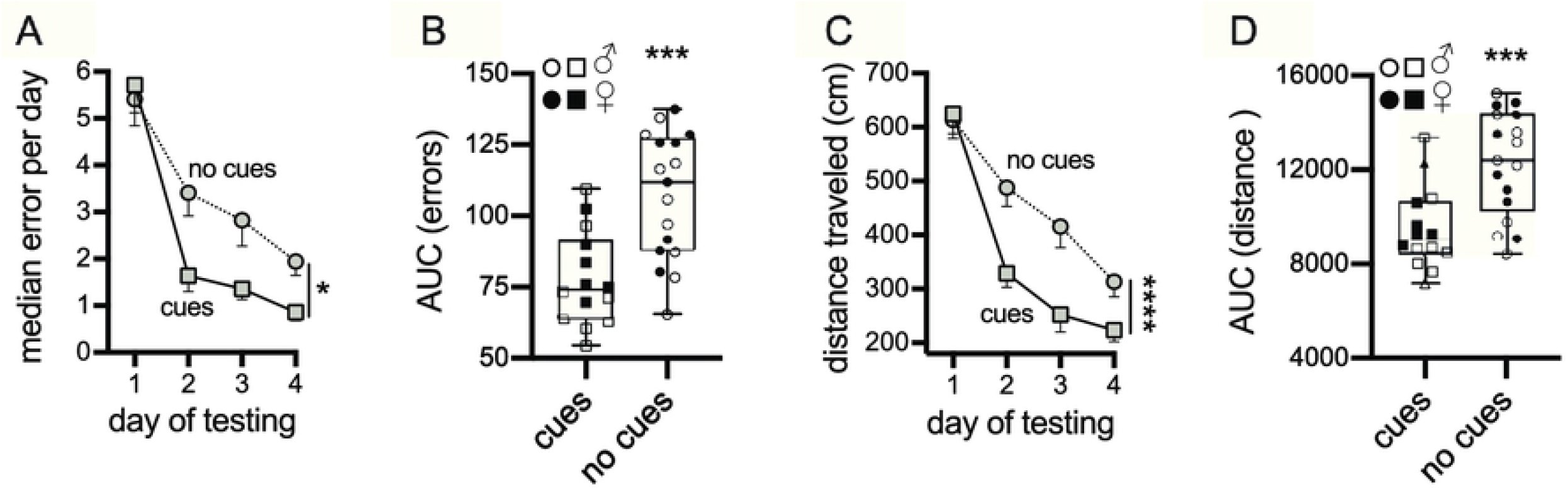
Evaluation of the use of cues and visible platform during RAWM test. Mice were exposed to a 7-trial test for a total of 4-day. **(A)** A learning curve of the median error per day is shown, a significant difference between mice using cues and mice not using cues was found (*p=0.0023). In **(B)** the area under the error curve confirmed that mice used cues to learn the task (***p=0.0003). Average (mean) distance **(C)** and area under the distance curve **(D)** showed that mice travelled more to locate the goal arm and escape platform as shown by the distance traveled (**C,** ****p<0.0001) and AUC **(D,** ***p=0.0007). In the AUC graphs male results are represented with the open symbol, instead females with closed symbol. Cues (n=14; 6/8 ♂/♀); no cues (n=17; 8/9 ♂/♀),

### Experiment 2: Scopolamine reduces learning ability in RAWM

Our next step was to use a pharmacological approach to validate this 7 trials RAWM protocol. Scopolamine is a “gold standard” non-selective muscarinic compound particularly used for validating learning and memory assay [31, 32]. These memory impairments are detected in a large variety of cognitive tasks: spontaneous alternation and novel spatial recognition Y-maze [31], Barnes maze test [33], contextual fear conditioning [33], MWM and RAWM [34]. Scopolamine is often used as a positive control compound for validating behavioral assays of learning and memory [35]. Scopolamine (1 mg/Kg) or vehicle were administered 30 minutes prior to trial 1 on each testing day. The median of errors made by scopolamine-treated mice was higher compared to vehicle-treated mice (Fig. 4 A, p<0.0001). Scopolamine-treated mice were able to learn the task, but slower compared to the vehicle group. AUC (errors x trials) is higher in scopolamine-treated mice as shown in Figure 4B (p<0.0001). Distance was also significantly different in scopolamine-treated mice (Fig 4 C and D, p<0.0001). Hyperactivity has been recorded after scopolamine treatment especially on day 1 as shown in Figure 4E, and this hyperactivity was reduced over the next three days of test. Representative heat maps are shown in Figure 4F. Vehicle-treated mice spent the majority of time in the goal arm as compared to the scopolamine-treated group that explored more of the maze across the 4-day test as demonstrated by a more yellow-red color across the entire RAWM apparatus.

**Figure 4:**
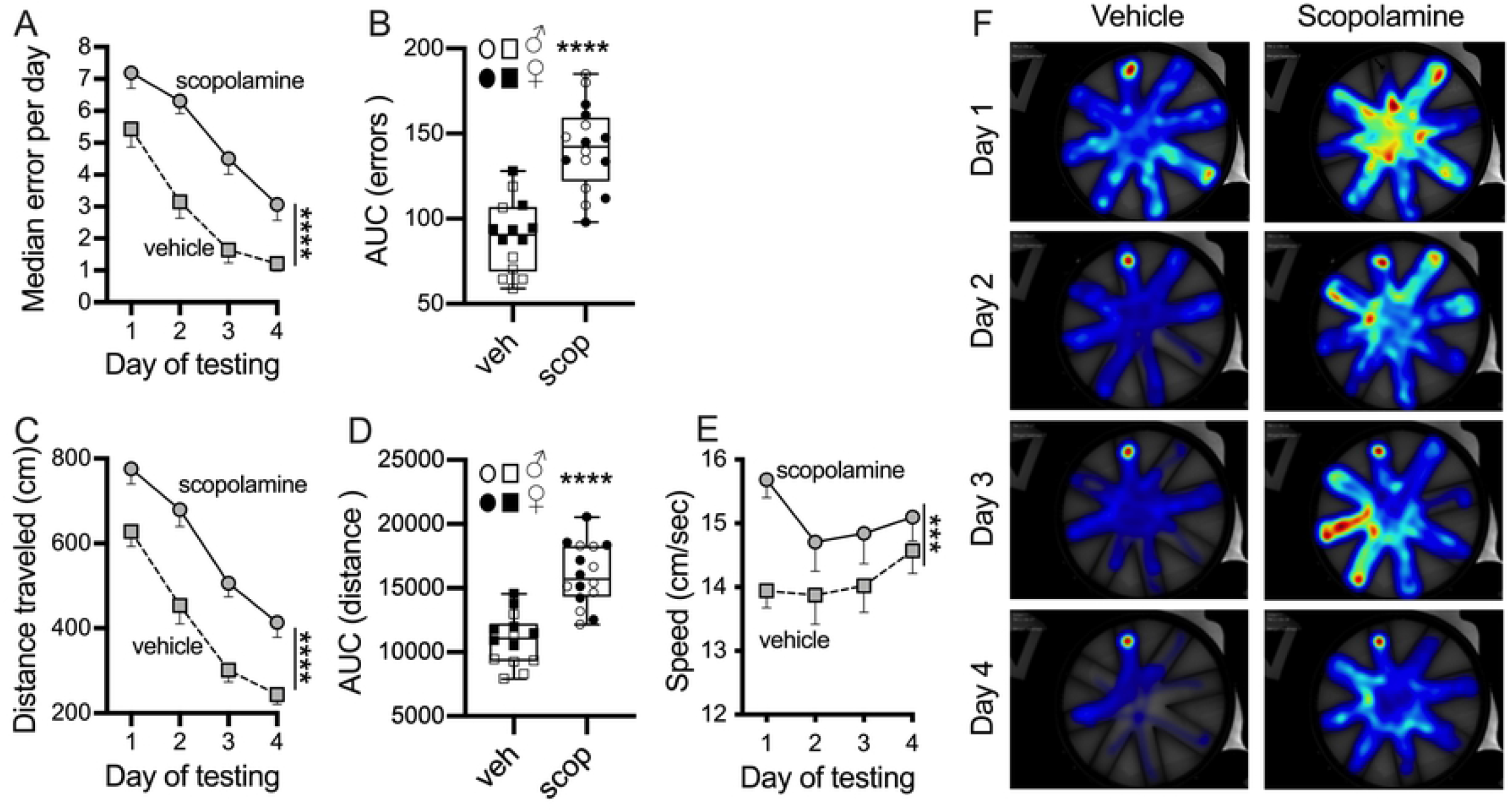
Validation of RAWM test using scopolamine as a positive control, to induce cognitive impairments. Male and female mice were tested in the RAWM test, scopolamine was given (1 mg/Kg; scop) 30 minutes prior to the start of testing on each day. **(A)** The median of errors made during the hidden platform over the 4-day test, and **(B)** area under the error curve showed that scopolamine disturbs learning and memory. In **(C)** the average distance traveled and **(D)** area under the distance curve confirmed that scopolamine-treated mice explore more the maze to escape. **(E)** Speed after scopolamine-treatment is increased. **(F)** Representation of time spent throughout the RAWM test using a heat map. Vehicle mice had a strong preference for the goal arm compared to scopolamine-treated mice that spend more time investigating the entire pool. Blue indicates that the mouse spends less time in that area of the maze, while red demarcates where the mouse spends the majority of its investigative time. In the AUC graphs male results are represented with the open symbol, instead females with closed symbol. ***p=0.0007, **** p<0.0001. Vehicle (n=14; 7/7 ♂/♀); scopolamine (n=16; 8/8 ♂/♀).

### Experiment 3: CHI impairs RAWM performance 2-week post-injury

In this experiment, we evaluated whether our 7-trial protocol was able to detect memory deficits in a mTBI model. In the past, it has been shown that the same animal model of CHI was memory impaired in a 6-arm RAWM when a 15-trial protocol and 2-day test [27, 36] and a 15-trial protocol and 4-day test [30] protocol were used. To compare differences between SHAM and CHI mice, we used the median of errors made during the 4-day test (Fig. 5 A). We found that sex had no effect on median errors per day (p=0.814) and mice that received a CHI performed worse on RAWM test than SHAM-operated mice (Fig. 5 A, p<0.0001). Both groups were able to learn the task over the 4-day test indicated by a reduction in the number of errors per day (Fig. 5 A, p<0.0001). Next, we compared the AUC for errors and similarly found that following CHI there was an increased AUC (error x trials) indicating more errors and a reduction in learning (Fig. 5 B, p=0.0002). Finally, when analyzing distance measures, we found that following CHI both distance travelled (Fig. 5 C, p<0.0001) and AUC (distance x trials) (Fig. 5 D, p=0.0001) were impaired.

**Figure 5:**
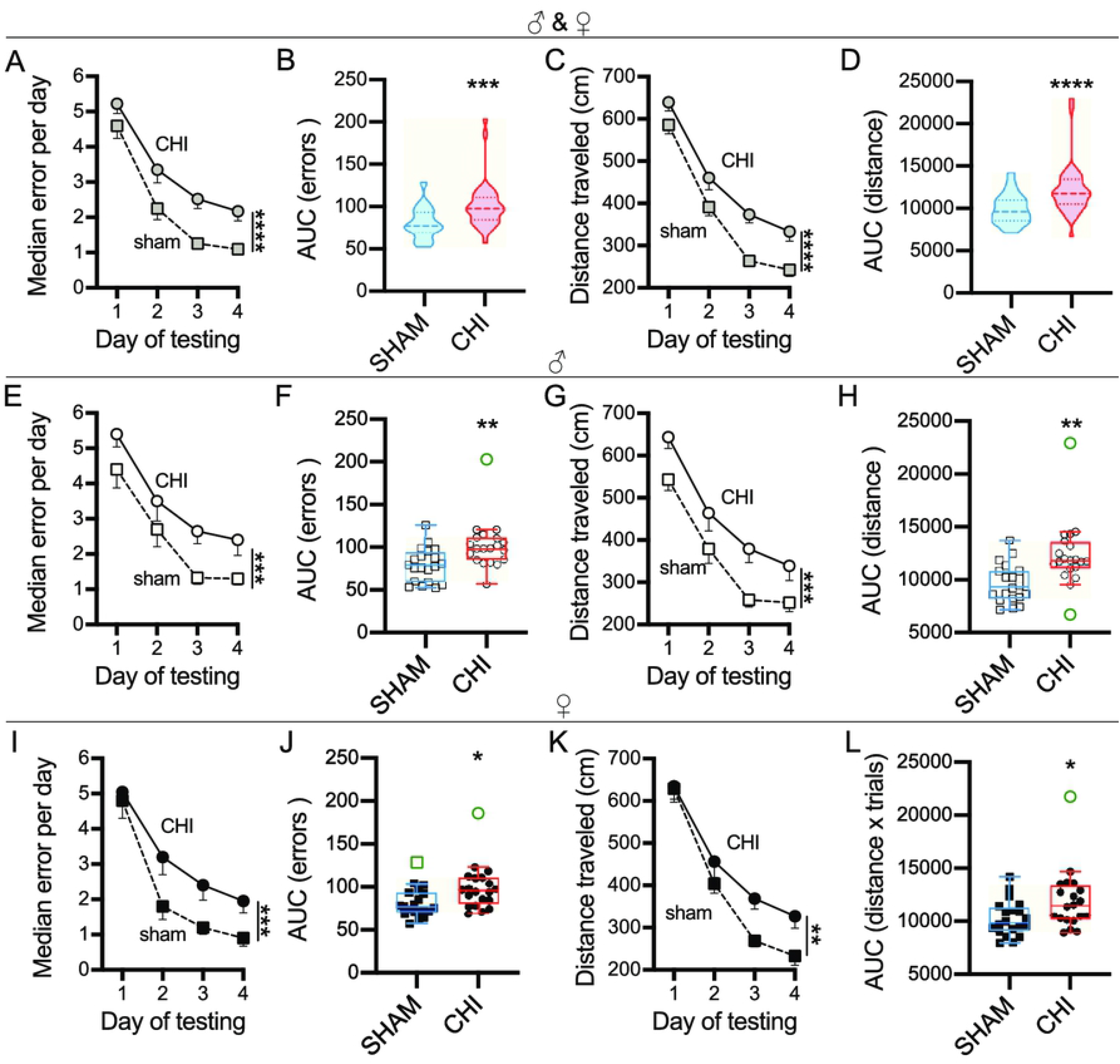
CHI induced memory deficits in RAWM at 2-week post-injury. Presented as males and females combined (first row), or males (second row) and females (third row) separately. **(A)** CHI- and SHAM-operated mice were able to learn the RAWM task, but CHI mice made more errors compared to SHAM mice (****p<0.0001). **(B**) Area under the error curve confirmed that CHI mice were memory impaired (*** p=0.0002). **(C)** Average distance travelled and **(D)** area under the distance curve show that CHI mice explored more than the SHAM group (****p<0.0001). Male mice following a CHI made more errors than the SHAM injured mice either by day **(E,** ***p=0.0004) or for the AUC **(F,** **p=0.0054). The trend held when data was plotted as distance traveled by day **(G,** ****p<0.0001) or for the AUC **(H,** **p=0.017). CHI was also found to cause worse performance in female mice for errors **(I,** ***p=0.0005; **J,** *p=0.016), and distance **(K,** **p=0.0014; L *p=0.026). Box-plot using Tukey method, with outliers shown in green. Sham (n=40; 20/20 ♂/♀); CHI (n=40; 20/20 ♂/♀).

## Discussion

Because of the heterogeneity of TBI, RAWM protocols can sometimes not be sensitive enough to detect injury-induced deficits in behavior. The goal of this paper was to provide a revised protocol for an 8-arm 4-day RAWM test. Our approach provided a sensitive protocol to detect cognitive impairments related to a single mild CHI at 2-week post injury. We believe that our protocol will be a useful resource for others attempting standardized behavioral assays, and those laboratories interested in detecting subtle differences in cognitive function in mouse models of CNS injury or disease.

A major finding of our study was that the reduction of number of trials significantly separates the acquisition curves of CHI and SHAM mice and the sex had no effect on results. Previously, we demonstrated that CHI mice had memory impairment in a 6-arm RAWM at 2 week post-CHI in a 4-day protocol and 15-trial per day [30] or in 2-day protocol and 15-trial per day [27, 36]. However, due to the knowledge that training schedule and spacing of training sessions can impact learning ability [10–13, 24, 25], we sought to determine if reduction of training increased the learning separation between mice affected by mTBI and controls. Our results confirmed that reduction of total number of trials increases the chance to separate learning impairment and this could be critical in a study where new compounds to reverse the injury effect are used [37] or when a smaller injury effect needs to be discovered [37]. A significant advantage using our new protocol is that more mice could be tested in one session, reducing time of testing and variability between groups.

Our results indicate that using a weaker training protocol, like reducing the number of trials, is more sensitive than a massive training to identify differences in cognition after a CHI [38]. Reduction in the number of training per day reduced fatigue and stress in the animals. Literature supports our idea that reducing the number of trials per day and increasing the inter-trial time helps the formation of long-term memory [10–12, 25, 26]. This is true not only in RAWM, but also in MWM and NOR test as well as in other memory tasks [14] indicating that our method may be able to be applied to other cognitive behavioral assays. Also, mice are less suitable to swimming compared to rats and are more sensitive to cold water [39], we think that exposing the mice to a limited number of trials per day will reduce the risk of hypothermia, fatigue and in particular the reduction of exposure to stress conditions [40].

While we believe that this protocol is more sensitive to detecting changes in mouse models, independent validation of behavioral protocols is important before conducting experiments. Each laboratory should conduct their own validation and include a group of mice to be used as positive control, in order to guarantee that the assay is reliable and consistent. Specially because it has been demonstrated that test standardization does not guarantee the same results across laboratories [41–43], due to variabilities that go beyond the scientist’s control. Laboratory environment, testing equipment, animal husbandry [41, 44], differences in the strains and genetic modification, experience [45] and sex of experimenters performing the test [46] play a robust role in behavioral results [45]. Validation is a key factor for obtaining trustable results and the protocol needs to be full of details to reduce the risk of unreproducible results across laboratory. A standard operating procedure should be created and used in each laboratory. Moreover, to enhance reproducibility an automatic scoring software, exclusion criteria and validation methods to train new experimenters should be established.

In conclusion, we demonstrated that: 1) the use of cues will improve the test acquisition, 2) reduction in the number of trials improves learning, and 3) a single mild CHI in mice could cause cognitive deficits detectable by the reduction of trials. We also provided a standard operating procedure and methods to validate the RAWM behavior for use in phenotyping other mouse models or when a pharmacological treatment would be considered.

## Acknowledgments

We would like to thank Collen N. Bondar, James B. Watson, and Henry C. Snider for their technical support and thoughtful comments on the manuscript.

## References

1. Barnes DE, Kaup A, Kirby KA, Byers AL, Diaz-Arrastia R, Yaffe K. Traumatic brain injury and risk of dementia in older veterans. Neurology. 2014;83(4):312–9. Epub 2014/06/27. doi: 10.1212/WNL.0000000000000616. PubMed PMID: 24966406; PubMed Central PMCID: PMCPMC4115602.

2. Johnson VE, Stewart W, Smith DH. Traumatic brain injury and amyloid-beta pathology: a link to Alzheimer’s disease? Nat Rev Neurosci. 2010;11(5):361–70. Epub 2010/03/11. doi: 10.1038/nrn2808. PubMed PMID: 20216546; PubMed Central PMCID: PMCPMC3979339.

3. Bodnar CN, Roberts KN, Higgins EK, Bachstetter AD. A Systematic Review of Closed Head Injury Models of Mild Traumatic Brain Injury in Mice and Rats. J Neurotrauma. 2019;36(11):1683–706. Epub 2019/01/22. doi: 10.1089/neu.2018.6127. PubMed PMID: 30661454; PubMed Central PMCID: PMCPMC6555186.

4. Buresova O, Bures J, Oitzl MS, Zahalka A. Radial maze in the water tank: an aversively motivated spatial working memory task. Physiol Behav. 1985;34(6):1003–5. Epub 1985/06/01. PubMed PMID: 4059369.

5. Hyde LA, Hoplight BJ, Denenberg VH. Water version of the radial-arm maze: learning in three inbred strains of mice. Brain Res. 1998;785(2):236–44. Epub 1998/06/06. PubMed PMID: 9518631.

6. Alamed J, Wilcock DM, Diamond DM, Gordon MN, Morgan D. Two-day radial-arm water maze learning and memory task; robust resolution of amyloid-related memory deficits in transgenic mice. Nat Protoc. 2006;1(4):1671–9. Epub 2007/05/10. doi: 10.1038/nprot.2006.275. PubMed PMID: 17487150.

7. Song S, Kong X, Acosta S, Sava V, Borlongan CV, Sanchez-Ramos J. Effects of an Inhibitor of Monocyte Recruitment on Recovery from Traumatic Brain Injury in Mice Treated with Granulocyte Colony-Stimulating Factor. Int J Mol Sci. 2017;18(7). Epub 2017/07/04. doi: 10.3390/ijms18071418. PubMed PMID: 28671601; PubMed Central PMCID: PMCPMC5535910.

8. Xiong XD, Xiong WD, Xiong SS, Chen GH. Age- and Gender-Based Differences in Nest-Building Behavior and Learning and Memory Performance Measured Using a Radial Six-Armed Water Maze in C57BL/6 Mice. Behav Neurol. 2018;2018:8728415. Epub 2018/06/02. doi: 10.1155/2018/8728415. PubMed PMID: 29854021; PubMed Central PMCID: PMCPMC5966705.

9. Kelley P, Whatson T. Making long-term memories in minutes: a spaced learning pattern from memory research in education. Front Hum Neurosci. 2013;7:589. Epub 2013/10/05. doi: 10.3389/fnhum.2013.00589. PubMed PMID: 24093012; PubMed Central PMCID: PMCPMC3782739.

10. Seese RR, Wang K, Yao YQ, Lynch G, Gall CM. Spaced training rescues memory and ERK1/2 signaling in fragile X syndrome model mice. Proc Natl Acad Sci U S A. 2014;111(47):16907–12. Epub 2014/11/12. doi: 10.1073/pnas.1413335111. PubMed PMID: 25385607; PubMed Central PMCID: PMCPMC4250145.

11. Kramar EA, Babayan AH, Gavin CF, Cox CD, Jafari M, Gall CM, et al. Synaptic evidence for the efficacy of spaced learning. Proc Natl Acad Sci U S A. 2012;109(13):5121–6. Epub 2012/03/14. doi: 10.1073/pnas.1120700109. PubMed PMID: 22411798; PubMed Central PMCID: PMCPMC3323981.

12. Kogan JH, Frankland PW, Blendy JA, Coblentz J, Marowitz Z, Schutz G, et al. Spaced training induces normal long-term memory in CREB mutant mice. Curr Biol. 1997;7(1):1–11. Epub 1997/01/01. PubMed PMID: 8999994.

13. Mandel RJ, Gage FH, Thal LJ. Enhanced detection of nucleus basalis magnocellularis lesion-induced spatial learning deficit in rats by modification of training regimen. Behav Brain Res. 1989;31(3):221–9. Epub 1989/01/01. PubMed PMID: 2914073.

14. Smolen P, Zhang Y, Byrne JH. The right time to learn: mechanisms and optimization of spaced learning. Nat Rev Neurosci. 2016;17(2):77–88. Epub 2016/01/26. doi: 10.1038/nrn.2015.18. PubMed PMID: 26806627; PubMed Central PMCID: PMCPMC5126970.

15. Anderson MJ, Jablonski SA, Klimas DB. Spaced initial stimulus familiarization enhances novelty preference in Long-Evans rats. Behav Processes. 2008;78(3):481–6. Epub 2008/03/25. doi: 10.1016/j.beproc.2008.02.005. PubMed PMID: 18358637.

16. Bello-Medina PC, Sanchez-Carrasco L, Gonzalez-Ornelas NR, Jeffery KJ, Ramirez-Amaya V. Differential effects of spaced vs. massed training in long-term object-identity and object-location recognition memory. Behav Brain Res. 2013;250:102–13. Epub 2013/05/07. doi: 10.1016/j.bbr.2013.04.047. PubMed PMID: 23644160.

17. Philips GT, Kopec AM, Carew TJ. Pattern and predictability in memory formation: from molecular mechanisms to clinical relevance. Neurobiol Learn Mem. 2013;105:117–24. Epub 2013/06/04. doi: 10.1016/j.nlm.2013.05.003. PubMed PMID: 23727358; PubMed Central PMCID: PMCPMC4020421.

18. Philips GT, Ye X, Kopec AM, Carew TJ. MAPK establishes a molecular context that defines effective training patterns for long-term memory formation. J Neurosci. 2013;33(17):7565–73. Epub 2013/04/26. doi: 10.1523/JNEUROSCI.5561-12.2013. PubMed PMID: 23616561; PubMed Central PMCID: PMCPMC3865502.

19. Wainwright ML, Zhang H, Byrne JH, Cleary LJ. Localized neuronal outgrowth induced by long-term sensitization training in aplysia. J Neurosci. 2002;22(10):4132–41. Epub 2002/05/23. doi: 20026347. PubMed PMID: 12019331.

20. Mauelshagen J, Sherff CM, Carew TJ. Differential induction of long-term synaptic facilitation by spaced and massed applications of serotonin at sensory neuron synapses of Aplysia californica. Learn Mem. 1998;5(3):246–56. Epub 1999/08/24. PubMed PMID: 10454368; PubMed Central PMCID: PMCPMC313806.

21. Beck CD, Schroeder B, Davis RL. Learning performance of normal and mutant Drosophila after repeated conditioning trials with discrete stimuli. J Neurosci. 2000;20(8):2944–53. Epub 2001/02/07. PubMed PMID: 10751447.

22. Menzel R, Manz G, Menzel R, Greggers U. Massed and spaced learning in honeybees: the role of CS, US, the intertrial interval, and the test interval. Learn Mem. 2001;8(4):198–208. Epub 2001/09/05. doi: 10.1101/lm.40001. PubMed PMID: 11533223; PubMed Central PMCID: PMCPMC311375.

23. Medin DL. The Comparative Study of Memory J Human Evol 1974;3:455–63

24. Light KR, Kolata S, Wass C, Denman-Brice A, Zagalsky R, Matzel LD. Working memory training promotes general cognitive abilities in genetically heterogeneous mice. Curr Biol. 2010;20(8):777–82. Epub 2010/03/30. doi: 10.1016/j.cub.2010.02.034. PubMed PMID: 20346673; PubMed Central PMCID: PMCPMC2910164.

25. Genoux D, Haditsch U, Knobloch M, Michalon A, Storm D, Mansuy IM. Protein phosphatase 1 is a molecular constraint on learning and memory. Nature. 2002;418(6901):970–5. Epub 2002/08/29. doi: 10.1038/nature00928. PubMed PMID: 12198546.

26. Zhang Y, Liu RY, Heberton GA, Smolen P, Baxter DA, Cleary LJ, et al. Computational design of enhanced learning protocols. Nat Neurosci. 2011;15(2):294–7. Epub 2011/12/27. doi: 10.1038/nn.2990. PubMed PMID: 22197829; PubMed Central PMCID: PMCPMC3267874.

27. Bachstetter AD, Webster SJ, Goulding DS, Morton JE, Watterson DM, Van Eldik LJ. Attenuation of traumatic brain injury-induced cognitive impairment in mice by targeting increased cytokine levels with a small molecule experimental therapeutic. J Neuroinflammation. 2015;12:69. Epub 2015/04/18. doi: 10.1186/s12974-015-0289-5. PubMed PMID: 25886256; PubMed Central PMCID: PMCPMC4396836.

28. Webster SJ, Van Eldik LJ, Watterson DM, Bachstetter AD. Closed head injury in an age-related Alzheimer mouse model leads to an altered neuroinflammatory response and persistent cognitive impairment. J Neurosci. 2015;35(16):6554–69. Epub 2015/04/24. doi: 10.1523/JNEUROSCI.0291-15.2015. PubMed PMID: 25904805; PubMed Central PMCID: PMCPMC4405562.

29. Chiarotti F. Statistical analysis of behavioral data. Curr Protoc Toxicol. 2005;Chapter 13:Unit 13 8. doi: 10.1002/0471140856.tx1308s25. PubMed PMID: 23045112.

30. Lyons DN, Vekaria H, Macheda T, Bakshi V, Powell DK, Gold BT, et al. A Mild Traumatic Brain Injury in Mice Produces Lasting Deficits in Brain Metabolism. J Neurotrauma. 2018;35(20):2435–47. Epub 2018/05/29. doi: 10.1089/neu.2018.5663. PubMed PMID: 29808778; PubMed Central PMCID: PMCPMC6196750.

31. Sukoff Rizzo SJ, Anderson LC, Green TL, McGarr T, Wells G, Winter SS. Assessing Healthspan and Lifespan Measures in Aging Mice: Optimization of Testing Protocols, Replicability, and Rater Reliability. Curr Protoc Mouse Biol. 2018;8(2):e45. Epub 2018/06/21. doi: 10.1002/cpmo.45. PubMed PMID: 29924918.

32. Klinkenberg I, Blokland A. The validity of scopolamine as a pharmacological model for cognitive impairment: a review of animal behavioral studies. Neurosci Biobehav Rev. 2010;34(8):1307–50. Epub 2010/04/20. doi: 10.1016/j.neubiorev.2010.04.001. PubMed PMID: 20398692.

33. Malikowska-Racia N, Podkowa A, Salat K. Phencyclidine and Scopolamine for Modeling Amnesia in Rodents: Direct Comparison with the Use of Barnes Maze Test and Contextual Fear Conditioning Test in Mice. Neurotox Res. 2018;34(3):431–41. doi: 10.1007/s12640-018-9901-7. PubMed PMID: 29680979; PubMed Central PMCID: PMCPMC6154175.

34. Salat K, Podkowa A, Mogilski S, Zareba P, Kulig K, Salat R, et al. The effect of GABA transporter 1 (GAT1) inhibitor, tiagabine, on scopolamine-induced memory impairments in mice. Pharmacol Rep. 2015;67(6):1155–62. doi: 10.1016/j.pharep.2015.04.018. PubMed PMID: 26481535.

35. Sukoff Rizzo SJ, Silverman JL. Methodological Considerations for Optimizing and Validating Behavioral Assays. Curr Protoc Mouse Biol. 2016;6(4):364–79. doi: 10.1002/cpmo.17. PubMed PMID: 27906464; PubMed Central PMCID: PMCPMC6054129.

36. Bachstetter AD, Morganti JM, Bodnar CN, Webster SJ, Higgins EK, Roberts KN, et al. The effects of mild closed head injuries on tauopathy and cognitive deficits in rodents: Primary results in wild type and rTg4510 mice, and a systematic review. Exp Neurol. 2020;326:113180. Epub 2020/01/14. doi: 10.1016/j.expneurol.2020.113180. PubMed PMID: 31930992.

37. Whiting MD, Kokiko-Cochran ON. Assessment of Cognitive Function in the Water Maze Task: Maximizing Data Collection and Analysis in Animal Models of Brain Injury. Methods Mol Biol. 2016;1462:553–71. Epub 2016/09/09. doi: 10.1007/978-1-4939-3816-2_30. PubMed PMID: 27604738.

38. Sebastian V, Diallo A, Ling DS, Serrano PA. Robust training attenuates TBI-induced deficits in reference and working memory on the radial 8-arm maze. Front Behav Neurosci. 2013;7:38. doi: 10.3389/fnbeh.2013.00038. PubMed PMID: 23653600; PubMed Central PMCID: PMCPMC3642509.

39. Bellush LL, Wright AM, Walker JP, Kopchick J, Colvin RA. Caloric restriction and spatial learning in old mice. Physiol Behav. 1996;60(2):541–7. Epub 1996/08/01. doi: 10.1016/s0031-9384(96)80029-1. PubMed PMID: 8840916.

40. Novais A, Monteiro S, Roque S, Correia-Neves M, Sousa N. How age, sex and genotype shape the stress response. Neurobiol Stress. 2017;6:44–56. Epub 2017/02/24. doi: 10.1016/j.ynstr.2016.11.004. PubMed PMID: 28229108; PubMed Central PMCID: PMCPMC5314441.

41. Wahlsten D, Metten P, Phillips TJ, Boehm SL, 2nd, Burkhart-Kasch S, Dorow J, et al. Different data from different labs: lessons from studies of gene-environment interaction. J Neurobiol. 2003;54(1):283–311. doi: 10.1002/neu.10173. PubMed PMID: 12486710.

42. Wahlsten D, Rustay NR, Metten P, Crabbe JC. In search of a better mouse test. Trends Neurosci. 2003;26(3):132–6. doi: 10.1016/S0166-2236(03)00033-X. PubMed PMID: 12591215.

43. Wahlsten D. Standardizing tests of mouse behavior: reasons, recommendations, and reality. Physiol Behav. 2001;73(5):695–704. doi: 10.1016/s0031-9384(01)00527-3. PubMed PMID: 11566204.

44. Crabbe JC, Wahlsten D, Dudek BC. Genetics of mouse behavior: interactions with laboratory environment. Science. 1999;284(5420):1670–2. Epub 1999/06/05. doi: 10.1126/science.284.5420.1670. PubMed PMID: 10356397.

45. Mandillo S, Tucci V, Holter SM, Meziane H, Banchaabouchi MA, Kallnik M, et al. Reliability, robustness, and reproducibility in mouse behavioral phenotyping: a cross-laboratory study. Physiol Genomics. 2008;34(3):243–55. Epub 2008/05/29. doi: 10.1152/physiolgenomics.90207.2008. PubMed PMID: 18505770; PubMed Central PMCID: PMCPMC2519962.

46. Sorge RE, Martin LJ, Isbester KA, Sotocinal SG, Rosen S, Tuttle AH, et al. Olfactory exposure to males, including men, causes stress and related analgesia in rodents. Nat Methods. 2014;11(6):629–32. Epub 2014/04/30. doi: 10.1038/nmeth.2935. PubMed PMID: 24776635.

